# phydms: Software for phylogenetic analyses informed by deep mutational scanning

**DOI:** 10.1101/121830

**Authors:** Sarah K. Hilton, Michael B Doud, Jesse D Bloom

## Abstract

**Background:** The evolution of protein-coding genes can be quantitatively modeled using phylogenetic methods. Recently, it has been shown that high-throughput experimental measurements of mutational effects made via deep mutational scanning can inform site-specific phylogenetic substitution models of gene evolution. However, there is currently no software tailored for such analyses.

**Results:** We describe software that efficiently performs phylogenetic analyses with substitution models informed by deep mutational scanning. This software, phydms, is ∼100-fold faster than existing programs that accommodate such substitution models. It can be used to compare the results of deep mutational scanning experiments to the selection on genes in nature. For instance, phydms enables rigorous comparison of how well different experiments on the same gene describe natural selection. It also enables the re-scaling of deep mutational scanning data to account for differences in the stringency of selection in the lab and nature. Finally, phydms can identify sites that are evolving differently in nature than expected from experiments in the lab.

**Conclusions:** The phydms software makes it easy to use phylogenetic substitution models informed by deep mutational scanning experiments. As data from such experiments becomes increasingly widespread, phydms will facilitate quantitative comparison of the experimental results to the actual selection pressures shaping evolution in nature.

## Background

It is widely appreciated that experiments in the lab can inform understanding of protein evolution in nature [1, 2]. Efforts to synthesize findings from experiments with evolutionary data have typically involved generating protein variants of interest, assaying their functionality in the lab, and qualitatively comparing the measured functionality of each variant to its evolutionary fate in nature [1, 2]. The recent advent of high-throughput deep mutational scanning techniques [3] has greatly expanded the potential of such research. For instance, numerous recent papers have reported measuring the effects of *all* amino-acid mutations on the functionality of a range of proteins [4, 5, 6, 7, 8, 9, 10, 11, 12, 13, 14, 15, 16, 17, 18, 19]. This flood of data necessitates new methods for comparing experimental measurements to evolution in nature, since simple qualitative inspection is insufficient when measurements are available for tens of thousands of mutants.

A solution is provided by the methods of molecular phylogenetics. Longstanding phylogenetic algorithms enable calculation of the statistical likelihood of an alignment of naturally occurring gene sequences given a phylogenetic tree and a model for the evolutionary substitution process [20, 21]. Deep mutational scanning data can be incorporated into this statistical framework via the substitution model [9]. Such experimentally informed codon models (ExpCM) of substitution can be used to test whether a deep mutational scanning experiment provides evolutionarily relevant information [9], compare the stringency of selection in nature and the lab [22, 23], assess how well different experiments describe natural selection on the same gene [12, 15], and identify sites that are evolving differently in nature than expected from experiments in the lab [23].

However, a hindrance to such analyses has been the lack of appropriate software. Prior work using ExpCM has involved re-purposing an existing software package (`HyPhy` [24] or `Bio++` [25]) to optimize the phylogenetic likelihood. Because these existing software packages are not designed for such site-specific models, the resulting analyses have been slow and cumbersome. Other software packages [26, 27, 28, 29] that handle site-specific codon substitution models are also not suitable, as they are designed to treat the effects of mutations as unknowns to be inferred rather than as values that have been measured *a priori*.

Here we describe `phydms`, software for <u>**phy**</u>logenetics informed by <u>**d**</u>eep <u>**m**</u>utational <u>**s**</u>canning. We show that `phydms` is ~100-fold faster than existing alternatives for performing analyses with ExpCM, and demonstrate how it can be used to quantitatively compare measurements from deep mutational scanning with selection on genes in nature. Readers who are interested in technical details of how `phydms` works should read the Implementation section; readers who are primarily interested in simply using `phydms` may prefer to jump directly to the Results and Discussion section.

## Implementation

### Substitution models

Here we briefly describe the codon substitution models implemented in `phydms`.

#### Experimentally informed codon models (ExpCM)

The basic ExpCM implemented in `phydms` are identical to those described in [23]. We recap these ExpCM to introduce nomenclature needed to understand the extensions described in the next few subsections.

In an ExpCM, rate of substitution ***P***_*r,xy*_ of site *r* from codon *x* to *y* is written in mutation-selection form [30, 31, 32] as 
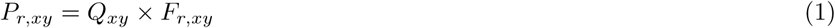
where *Q*_*xy*_ is proportional to the rate of mutation from *x* to *y*, and ***F***_*r,xy*_ is proportional to the probability that this mutation fixes. The rate of mutation ***Q***_*xy*_ is assumed to be uniform across sites, and takes an HKY85-like [33] form: 
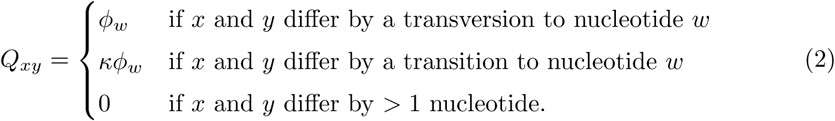

The *K* parameter represents the transition-transversion ratio, and the *ϕ*_*w*_ values give the expected frequency of nucleotide *w* in the absence of selection on amino-acid substitutions, and are constrained by 1 = Σ_*w*_*ϕ*_*w*_.

The deep mutational scanning data are incorporated into the ExpCM via the *F*_*r,xy*_ terms. The experiments measure the preference *π*_*r,a*_ of every site *r* for every amino-acid *a* (see the **Results and Discussion** section for more details on these preferences). The *F*_*r,xy*_ terms are defined in terms of these experimentally measured amino-acid preferences as 
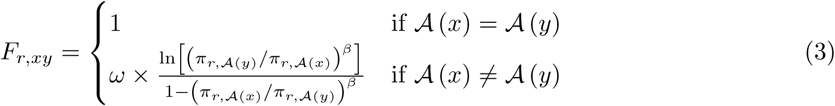
where 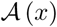 is the amino-acid encoded by codon *x, β* is the stringency parameter, and *ω* is the relative rate of nonsynonymous to synonymous substitutions after accounting for the amino-acid preferences. As shown in Figure 1, Equation 3 implies that mutations to more preferred amino acids are favored, and mutations to less preferred amino acids are disfavored. The functional form in Equation 3 was derived by Halpern and Bruno [30] and under certain (probably unrealistic) population-genetic assumptions; under these assumptions, *β* is related to the effective population size. When *β* > 1, natural evolution favors the same mutations as the experiments but with greater stringency. The ExpCM have six free parameters (three *ϕ*_*w*_ values, *κ, β*, and *ω*). The preferences *π*_*r,a*_ are *not* free parameters since they are determined by an experiment independent of the sequence alignment being analyzed.

**Figure 1.**
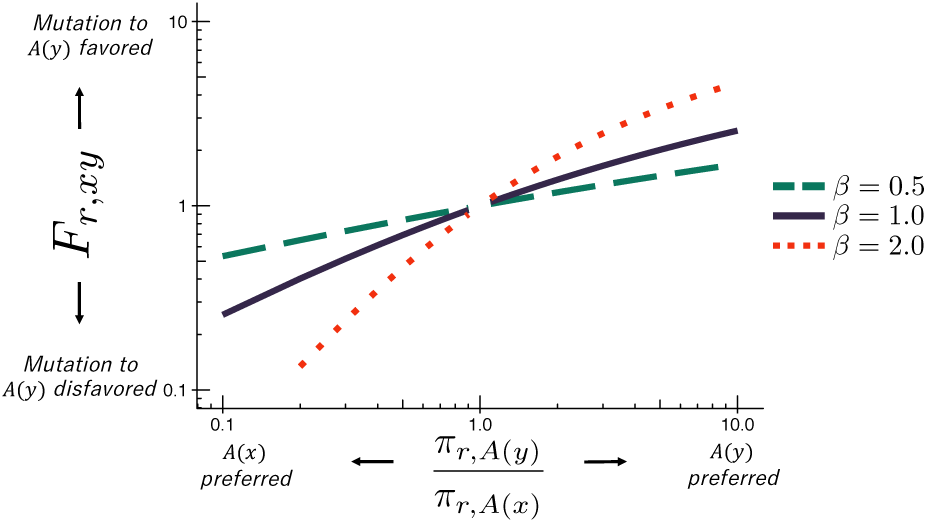
The ExpCM fixation term *F*_*r,xy*_. In an ExpCM, the rate of fixation of a mutation from codon *x* to codon *y* depends on the experimentally measured preferences of the amino acids *A*(*x*) and *A*(*y*) encoded by these codons. Mutations to preferred amino acids, with 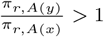, result in a larger *F*_*r,xy*_, and so are anticipated to fix more often. Re-scaling the preferences by a stringency parameter *β* ≠ 1 to reflect differences in selection between the lab and nature modulates *F*_*r,xy*_. When β > 1, the selection for preferred amino acids is exaggerated. When β < 1, the selection for preferred amino acids is attenuated.

#### ExpCM with empirical nucleotide frequency parameters

Phylogenetic substitution models commonly set the nucleotide frequency parameters (*ϕ*_*w*_ in the case of an ExpCM) so that the model’s stationary state equals the empirical frequencies of the characters in the alignment. Setting the frequency parameters in this way reduces the number of parameters that must be optimized by maximum likelihood. Empirically setting the nucleotide frequency parameters is easy for substitution models where the stationary state only depends on these parameters.

However, the situation for an ExpCM is more complex. The *ϕ*_*w*_ values give the expected nucleotide frequencies in the *absence* of selection on amino acids, but in an ExpCM there is site-specific selection on amino acids. Therefore, the stationary state of an ExpCM also depends on other quantities: the stationary state frequency *p*_*r,x*_ of codon *x* at site *r* is [23] 
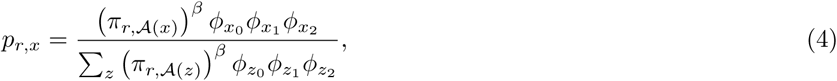
where *x*_*k*_ indicates the nucleotide at position *k* in codon *x*. As this equation makes clear, the stationary state of an ExpCM depends on the preferences *π*_*r,a*_ and stringency parameter *β* as well as the nucleotide frequency parameters *ϕ*_*w*_.

So for an ExpCM, setting *ϕ*_*w*_ empirically means choosing their values such that the alignment frequency *g*_*w*_ of nucleotide *w* is as expected given the stationary state *p*_*r,x*_. This will be the case if the following equation holds for all *w*: 
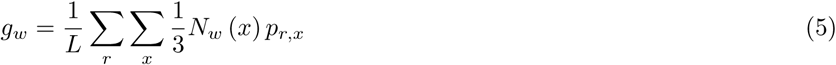
where *L* is the length of the gene in codons, *r* ranges over all codon sites, *x* ranges over all codon identities, and *N*_*w*_(*x*) is the number of occurrences of nucleotide *w* in codon *x*. We could not analytically solve this system of equations for *ϕ*_*w*_ in terms of *g*_*w*_, so we instead used a non-linear equation solver to determine the values as detailed in Additional file 1. Calculating *ϕ*_*w*_ empirically in this fashion is the default for `phydms`. If you instead want to fit the *ϕ*_*w*_ values, use the `−−fitphi` option.

#### ExpCM with gamma-distributed nonsynonymous-to-synonymous rate parameter

A common extension to traditional non-site-specific codon substitution models is to allow the dN/dS ratio ω to come from several discrete categories by making the overall likelihood at each site a linear combination of the likelihood computed for each category [34, 35]. Such models are not site-specific since sites are not assigned to a category during likelihood optimization, but they do capture the idea that there are different strengths of selection on nonsynonymous mutations across sites.

One variant of this approach draws *ω* from a discrete gamma distribution. This variant is referred to as the M5 variant [35] in `PAML` [36]. We implemented a similar approach for ExpCM, following [37] to draw the *ω* in Equation 3 from the means of equally weighted gamma-distributed categories (by default there are four categories). This option can be used via the `−−gammaomega` switch to `phydms`, and adds one free parameter, since there are two parameters controlling the gamma distribution (a shape and inverse-scale parameter) rather than a single *ω*. This option increases the runtime by ~5-fold.

Using a gamma-distributed ω typically leads to less of an improvement in fit for ExpCM than for non-site-specific models, since much of the site-to-site variation in the selection is already captured by the amino-acid preferences. However, it can still lead to substantial improvements if a subset of sites are under diversifying selection or if the preferences do not fully capture selection on nonsynonymous mutations.

#### Traditional YNGKP-style models

To enable comparison of ExpCM with non-site-specific substitution models, `phydms` implements several of these more traditional models. These models are referred to as YNGKP as they are variants of the Goldman-Yang style models described by Yang, Nielsen, Goldman, and Krabbe-Pedersen [35]. The M0 and M5 YNGKP models are implemented in `phydms`. The M0 variant optimizes a single dN/dS ratio (*ω*) and so is comparable with the basic ExpCM, while the M5 variant draws *ω* from a gamma distribution and so is comparable to ExpCM with the `−−gammaomega` option. The equilibrium codon frequencies are calculated empirically after correcting for stop codons as described in [38] (the so-called CF3X4 method). The M0 variant has 11 parameters (9 empirical nucleotide frequencies plus *ω* and *κ*), while the M5 variant has 12 parameters (there are two parameters for the gamma distribution over *ω*).

YNGKP models are less computationally expensive than ExpCM since they are not site-specific. Therefore, YNGKP models are faster than ExpCM in `phydms`. However, `phydms` is *not* optimized for maximal speed with YNGKP models, so if you are only using those models then consider using `PAML` [36] or `HyPhy` [24].

### Gradient-based optimization of the likelihood

Given one of the substitution models described above and a fixed phylogenetic tree topology, `phydms` numerically optimizes the model parameters and branch lengths to their maximum likelihood values via the Felsenstein pruning algorithm [21]. Numerical optimization generally requires fewer steps if the gradient of the objective function with respect to free parameters is computed explicitly [39], although this advantage can be offset by the cost of computing the gradient. We were unable to find clear published comparisons of the efficiency of phylogenetic optimization with and without an explicit gradient, although there is literature describing how the gradient (and Hessian matrix of second derivatives) can be computed [40].

We chose to use gradient-based optimization for `phydms` under the supposition that it might be more efficient. The first derivatives with respect to branch lengths and virtually all the model parameters can be computed analytically, propagated through the matrix exponentials using the formula provided by [41] (see also [40, 42]), and evaluated along the tree by applying the chain rule to the Felsenstein pruning algorithm. For the ExpCM empirical nucleotide frequencies *ϕ*_*w*_ and the gamma-distributed *ω*, we used the numerical finite-difference method to compute small portions of the derivatives for which we could not derive analytic results. Additional file 1 details how `phydms` computes the likelihood and its gradient.

For the optimization, we used the limited-memory BFGS optimizer with bounds [43, 44, 45]. This optimizer uses the gradient, although this can be turned off with the `−−nograd` option to `phydms` (doing so is *not* recommended as the accuracy of `phydms` without gradients has not been extensively tested). Rather than optimizing model parameters and branch lengths simultaneously, `phydms` takes an iterative approach. First the model parameters are simultaneously optimized along with a single scaling parameter that multiplies all branch lengths. After this optimization has converged, all branch lengths are simultaneously optimized while holding the model parameters constant. This process is repeated until further optimization leads to negligible improvement in the likelihood. Note that simultaneous optimization of all branch lengths appears to be the minority approach in phylogenetics software [46] and is stated to be less efficient than one-at-a-time optimization in [47]; however, we found it to work effectively on the trees that we tested. The rationale for iterating between model parameters and branch lengths is that optimization of the former tends to be more difficult and more costly in terms of the gradient computation. If you simply want to scale branch lengths by a single parameter rather than optimize them, you can use the `−−brlen scale` option. In other contexts, scaling but not individually optimizing branch lengths has been shown to reduce runtime with little effect on final model parameters if the initial tree is reasonably accurate [47, 48].

### Design and implementation of phydms

The `phydms` software is written in `Python`. Most of the numerical computation is performed with `numpy` and `scipy`, and a few parts of the code are written in compiled `C` extensions created via `cython`. The limited-memory BFGS optimizer used by `phydms` is the one provided with `scipy.optimize`. The most computationally costly part of the optimization performed by `phydms` is the matrix-matrix multiplication performed when computing exponentials of the transition matrix, and the second most costly part is the matrix-vector multiplication performed while implementing the Felsenstein pruning algorithm. Both these steps are performed using `BLAS` subroutines called via `scipy`.

In addition to the core `phydms` program, the software is distributed with auxillary programs that make it easy to prepare alignments (`phydms_prepalignment`) and run multiple models for comparison (`phydms_comprehensive`). Importantly, `phydms` currently does *not* infer phylogenetic tree topologies, but rather optimizes branch lengths and model parameters given a topology. The tree topology must therefore be inferred using another program such as `RAxML` [49] with a simpler substitution model.

### Visualization of the results with logoplots

It is often instructive to visualize the amino-acid preferences that are used to inform ExpCM, as these preferences determine the unique properties of the models. In addition, visualization can help understand how the stringency parameter *β* optimized by `phydms` re-scales the preferences to increase concordance with natural selection. To aid such visualizations, `phydms` comes with an auxiliary program (`phydms_logoplot`) that renders the amino-acid preferences in the form of logoplots via the `weblogo` libraries [50]. The Results and Discussion section below shows example logoplots.

## Results and Discussion

### Testing phydms on two different genes

In the sections below, we use **phydms** to compare deep mutational scanning measurements to natural sequence evolution for two genes: influenza hemagglutinin (HA) and *β*-lactamase. We choose these genes because there are multiple published deep mutational scanning datasets for each.

Analysis with ExpCM requires three pieces of input data: the experimentally measured amino-acid preferences, an alignment of naturally occurring gene sequences, and a phylogenetic tree topology. The tree topology can be inferred from the sequence alignment. But like most other software for codon-based phylogenetic analyses [24, 36], `phydms` is not designed to infer the tree topology – instead, it provides easy ways to infer this tree using other software such as `RAxML` [49].

To prepare the required input data, we followed the workflow in Figure 2. The deep mutational scanning experiments on HA [10, 15] directly reported amino-acid preferences. However, the two *β*-lactamase deep mutational scanning experiments [6, 11] reported enrichment ratios for each mutation rather than amino-acid preferences. There is a simple relationship between enrichment ratios and amino-acid preferences: the preferences are the enrichment ratios after normalizing the values to sum to one at each site, enabling easy conversion between the two data representations (Figure 2).

**Figure 2.**
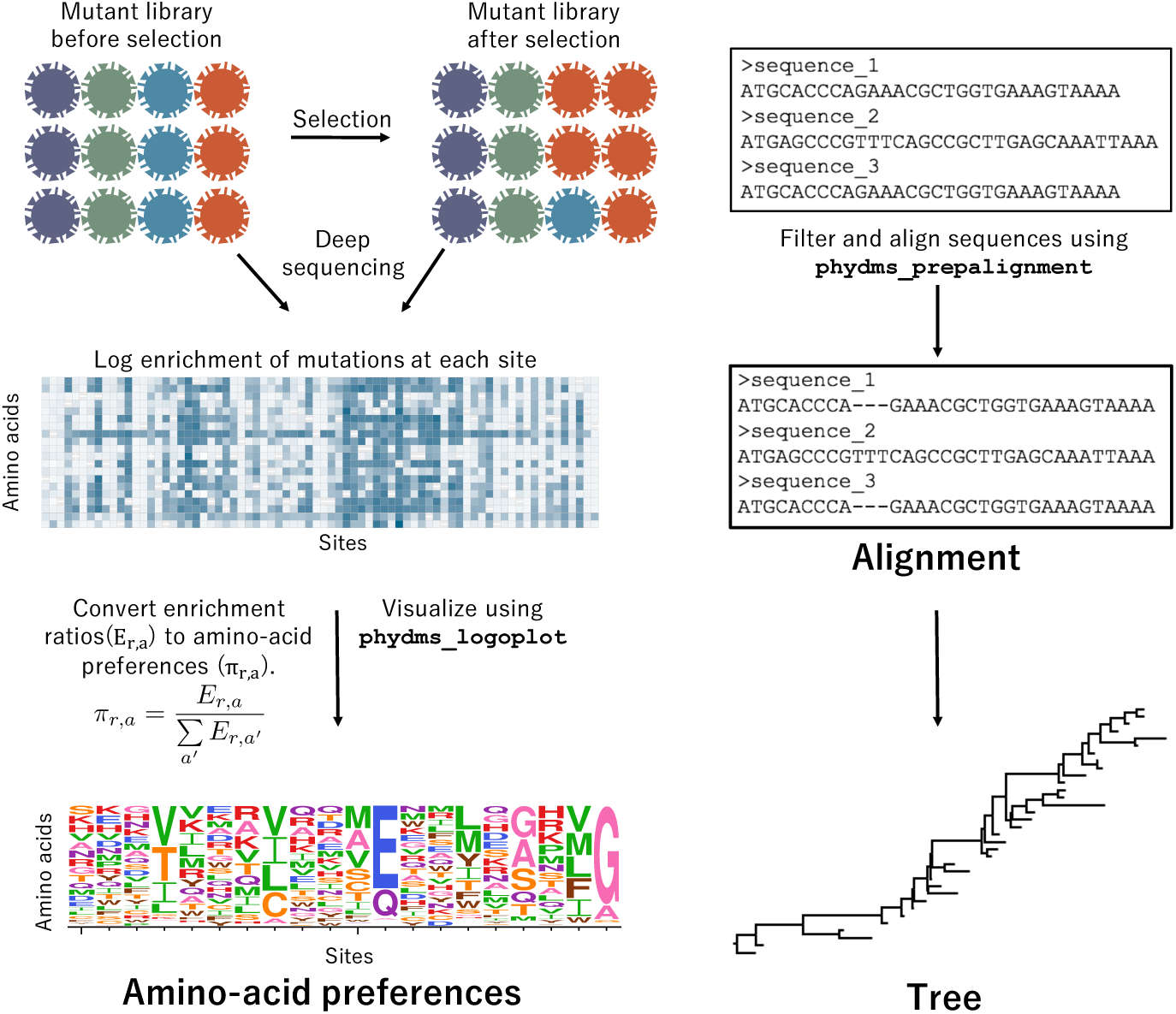
Workflow for preparing input data to phydms. Analysis with `phydms` requires amino-acid preferences measured by deep mutational scanning, a codon-level alignment of naturally occurring sequences, and a phylogenetic tree topology. Deep mutational scanning involves performing a functional selection on a library of mutant genes, and using deep sequencing to quantify the enrichment or depletion of each mutation after selection. To process the deep mutational scanning data referred to in Table 1, we converted enrichment ratios into amino-acid preferences by normalizing the values to sum to one at each site. We created a filtered, codon-level alignment of naturally occurring sequences using `phydms_prepalignment`. We used `phydms_comprehensive` to automatically generate a tree topology from the filtered alignment using `RAxML`.

We also created codon-level alignments of naturally occurring HA and *β*-lactamase sequences using `phydms_prepalignment`. The alignments were trimmed to contain only sites for which amino-acid preferences were experimentally measured. Table 1 summarizes basic information about these alignments.

**Table 1.** Alignments and deep mutational scanning (DMS) studies for HA and *β*-lactamase.

| gene | DMS studies | residues in protein | residues with DMS data | number of sequences |
| --- | --- | --- | --- | --- |
| HA | [15], [10] | 565 | 564 | 34 |
| β-lactamase | [11], [6] | 285 | 263 | 50 |

### Test if deep mutational scanning is informative about natural selection

A first simple test is whether the deep mutational scanning experiment provides any information that is relevant to natural selection on the gene in question. This can be determined by testing whether an ExpCM that uses the experimental data outperforms a standard substitution model that is agnostic to the site-specific preferences measured in the experiments.

To perform such a test, we used `phydms_comprehensive` to fit several substitution models to the alignment of HA sequences. This program automatically generates a phylogenetic tree topology from the alignment using `RAxML` [49]. It then fits an ExpCM (in this case informed by the deep mutational scanning data in Doud and Bloom [15]) as well as several substitution models that do not utilize site-specific experimental information. The analysis was performed by running the following command on the input data in Additional file 2:

~~~
phydms_comprehensive results/ HA_alignment.fasta HA_Doud_prefs.csv ‐‐raxml raxml
~~~

Table 2 lists the four tested substitution models: the ExpCM, an ExpCM with the amino-acid preferences averaged across sites, and the M0 and M5 variants of the standard Goldman-Yang style substitution models [35]. The ExpCM with averaged preferences is a sensible control because averaging the preferences eliminates any experimental information specific to individual sites in the protein. Because the models have different numbers of free parameters, they are best compared using Akaike Information Criterion (AIC) [51], which compares log likelihoods after correcting for the number of free parameters. Table 2 shows that the ExpCM has a much smaller AIC than the other models (ΔAIC > 2000 for all of the other models). This result shows that the experimentally measured amino-acid preferences contain information about natural selection on HA, since a substitution model informed by these preferences greatly outperforms generic substitution models that do not utilize experimental information.

**Table 2.** Fitting of ExpCM informed by the HA preferences measured in [15] to natural sequences using `phydms_comprehensive`. Full code, data, and results are in Additional file 2.

| model | ΔAIC | log likelihood | number of parameters | parameter values |
| --- | --- | --- | --- | --- |
| ExpCM with Doud preferences | 0.0 | -4877.7 | 6 | $\beta=2.11, \kappa=5.14, \omega=0.52$ |
| ExpCM with averaged Doud preferences | 2090.6 | -5922.9 | 6 | $\beta=0.68, \kappa=5.36, \omega=0.22$ |
| YNGKP_M5 | 2113.5 | -5928.4 | 12 | $\alpha_\omega=0.30, \beta_\omega=1.42, \kappa=4.68$ |
| YNGKP_M0 | 2219.6 | -5982.5 | 11 | $\kappa=4.61, \omega=0.20$ |

### Re-scale deep mutational scanning data to stringency of natural selection

Even if a deep mutational scanning experiment measures the authentic natural selection on a gene, the stringency of selection in the experiment is not expected to match the stringency of selection in nature. Differences in the stringency of selection can be captured by the ExpCM stringency parameter *β*. If selection in nature prefers the same amino acids as the selection in lab but with greater stringency, *β* will be fit to a value > 1. Conversely, if selection in nature does not prefer the lab-favored mutations with as much stringency as the deep mutational scan, *β* will be fit to a value < 1. Table 2 shows that ExpCM for HA informed by the experiments in [15] have *β* = 2.11, indicating that natural selection favors the experimentally preferred amino acids with higher stringency than selection in the lab.

The effect of this stringency re-scaling of the preferences can be visualized using `phydms_logoplot` as shown in Figure 3. Re-scaling by the optimal stringency parameter of 2.11 exaggerates the selection for experimentally preferred amino acids. Conversely, if the analysis had fit a stringency parameter < 1, this would have flattened the experimental measurements, and when *β* = 0 all information from the experiments is lost (Figure 3). Because selection in the lab can probably never be tuned to exactly match that in nature, stringency re-scaling is a valuable method to standardize measurements across experiments.

**Figure 3.**
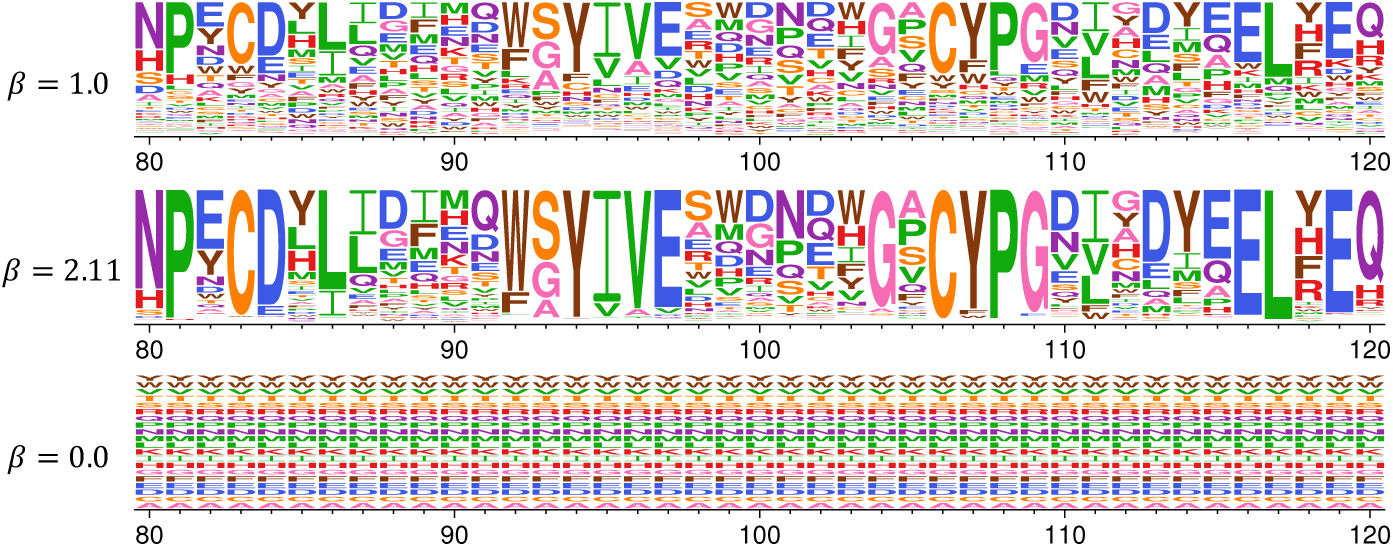
Re-scaling of amino-acid preferences to reflect the stringency of selection in nature. Analysis with `phydms` optimizes a stringency parameter *β* that relates the stringency of selection for preferred amino acids in the deep mutational scanning experiment to that in nature. When *β* = 1, the preferences are same as the values measured in the lab, suggesting a similar stringency of selection in the experiments and in nature. When *β* > 1, selection in nature prefers the same amino acids as selection in lab but with greater stringency – therefore, highly preferred amino acids increase in preference while lowly preferred amino acids decrease. When *β* < 1, selection in nature has less preference than the experiments for mutations favored in the lab, and when β = 0 then all site-specific information is lost. The actual optimized stringency parameter reported in Table 2 is β = 2.11. We generated the logoplots shown above from the input data in Additional file 3 with the following commands:

~~~
phydms_logoplot HA_Doud_1.pdf ‐‐prefs HA_Doud_prefs_short.csv
phydms_logoplot HA_Doud_2_11.pdf ‐‐prefs HA_Doud_prefs_short.csv ‐‐stringency
     2.11
phydms_logoplot HA_Doud_0.pdf ‐‐prefs HA_Doud_prefs_short.csv ‐‐stringency 0
~~~

### Compare how well different experiments capture natural selection

The amino-acid preferences for both HA and *β*-lactamase have each been measured by two independent experiments. For each gene, which of these experiments better captures natural selection?

We can address this question by comparing ExpCM informed by each experiment. For *β*-lactamase, this means comparing the preferences measured by Stiffler et al [11] to those measured by Firnberg et al [6]. We did this with `phydms_comprehensive` by running the following command on the input data in Additional file 4:

~~~
phydms_comprehensive results/ betaLactamase_alignment.fasta
     betaLactamase_Stiffler_prefs.txt betaLactamase_Firnberg_prefs.txt ‐‐raxml raxml
~~~

Table 3 shows that the ExpCM informed by the data of Stiffler et al [11] outperform ExpCM informed by the data of Firnberg et al [6], with a ΔAIC of 96.2. Therefore, the former experiment better reflects natural selection on *β*-lactamase. However, both experiments are clearly informative, as they both greatly outperform the traditional YNGKP models.

**Table 3.** Comparison of multiple *β*-lactamase deep mutational scanning results using phydms_comprehensive. Full code, data, and results are in Additional file 4.

| model | ΔAIC | log likelihood | number of parameters | parameter values |
| --- | --- | --- | --- | --- |
| ExpCM with Stiffler preferences | 0.0 | -2581.3 | 6 | $\beta=1.31, \kappa=2.67, \omega=0.72$ |
| ExpCM with Firnberg preferences | 96.2 | -2629.4 | 6 | $\beta=2.42, \kappa=2.60, \omega=0.63$ |
| YNGKP_M5 | 739.2 | -2944.9 | 12 | $\alpha_\omega=0.30, \beta_\omega=0.49, \kappa=3.02$ |
| YNGKP_M0 | 841.0 | -2996.8 | 11 | $\kappa=2.39, \omega=0.28$ |

We made a similar comparison of the two deep mutational scans of HA. As summarized in Table 4 (and detailed in Additional file 5), the deep mutational scanning of Doud and Bloom [15] better describes the natural evolution than the experiments of Thyagarajan and Bloom [10] (ΔAIC of 44.2). Again, both experiments are clearly informative, as they both greatly outperform the YNGKP models.

**Table 4.** Comparison of multiple HA deep mutational scanning results using phydms_comprehensive. Full code, data, and results are in Additional file 5.

| model | ΔAIC | log likelihood | number of parameters | parameter values |
| --- | --- | --- | --- | --- |
| ExpCM with Doud preferences | 0.0 | -4877.7 | 6 | $\beta=2.11, \kappa=5.14, \omega=0.52$ |
| ExpCM with Thyagarajan preferences | 44.2 | -4899.7 | 6 | $\beta=1.72, \kappa=4.94, \omega=0.55$ |
| YNGKP_M5 | 2113.5 | -5928.4 | 12 | $\alpha_\omega=0.30, \beta_\omega=1.42, \kappa=4.68$ |
| YNGKP_M0 | 2219.6 | -5982.5 | 11 | $\kappa=4.61, \omega=0.20$ |

Overall, these results show how `phydms` can rigorously compare how well different experiments describe the evolution of a gene.

### Identify sites of diversifying selection

In some cases, a few sites may evolve differently in nature than expected from the experiments in the lab. For instance, sites under diversifying selection for amino-acid change will experience more nonsynonymous substitutions than expected given the experimentally measured amino-acid preferences. Such sites can be identified by using the `−−omegabysite` option to fit a parameter *ω*_*r*_ that gives the relative rate of nonsynonymous to synonymous substitutions after accounting for the experimentally measured preferences for each site *r* [23]. If the preferences capture all the selection on amino acids, then we expect *ω*_*r*_ = 1. Sites with *ω*_*r*_ > 1 are under diversifying selection for amino-acid change, while sites with *ω*_*r*_ < 1 are under additional purifying selection not measured in the lab.

We tested for diversifying selection in HA by running the following command on the data in Additional file 6:

~~~
phydms HA_alignment.fasta HA_RAxML_tree.newick ExpCM_HA_Doud_prefs.csv results/
     ‐‐omegabysite
~~~

The results are visualized in Figure 4. While the majority of sites are evolving with *ω*_*r*_ not significantly different from one, some sites show evidence of *ω*_*r*_ > 1. As described in [23], these sites may be under diversifying selection due to immune pressure. Overall, this analysis shows how `phydms` can identify individual sites evolving differently in nature than expected from experiments in the lab.

**Figure 4.**
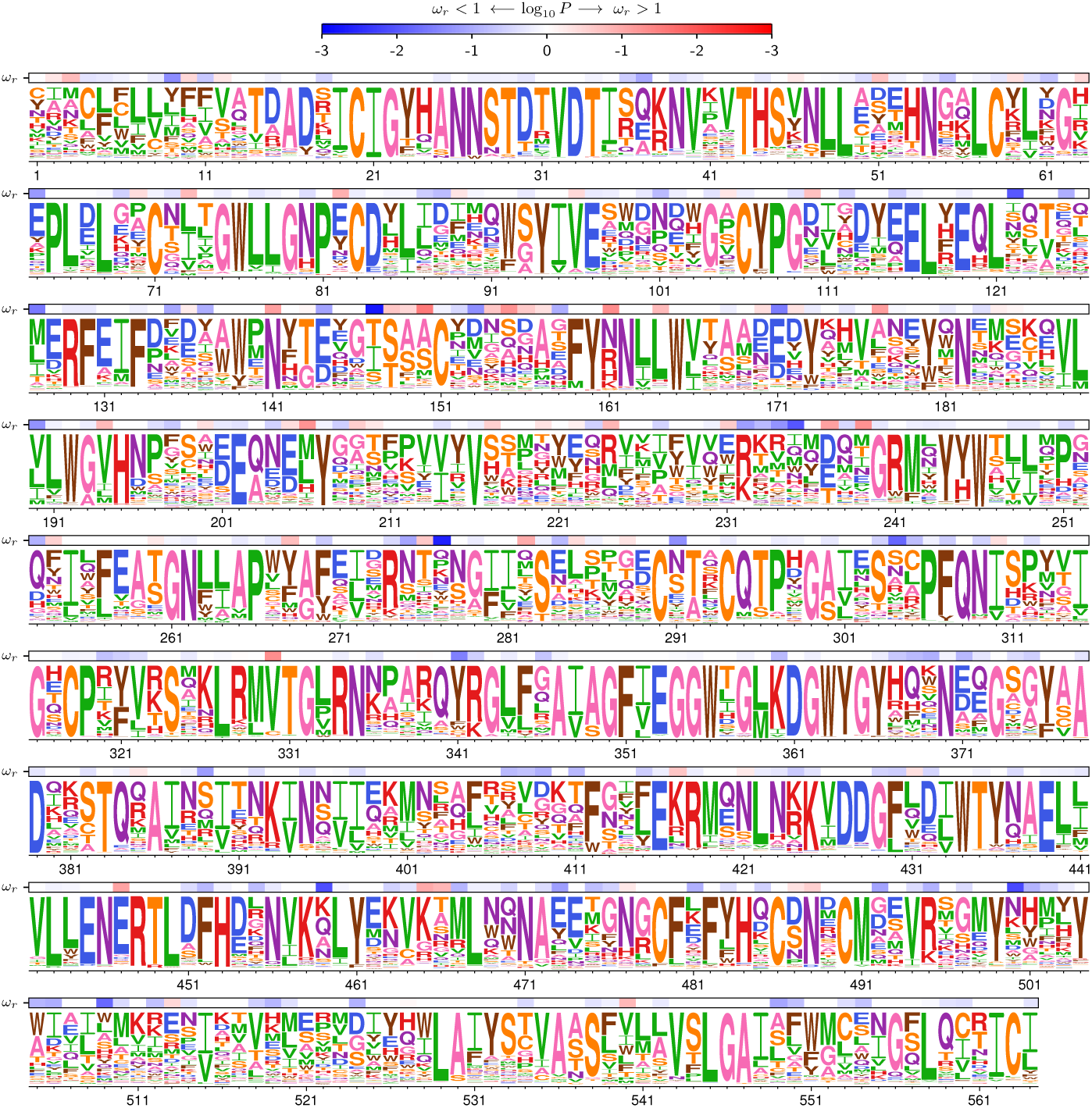
Identifying sites of diversifying selection. The `phydms` option `−−omegabysite` fits a site-specific value for *ω*_*r*_, which gives the relative rate of nonsynonymous to synonymous substitutions at site *r* after accounting for the selection due to the amino-acid preferences. This figure shows the results of such an analysis for HA. The overlay bar represents the strength of evidence for *ω*_*r*_ being greater (red) or less (blue) than one. Because this approach accounts for the constraints due to the amino-acid preferences, it can identify sites evolving faster than expected even if their absolute relative rates of nonysnonymous to synonymous substitutions do not significantly differ from one [23]. The logoplot in this figure uses the stringency parameter value of *β* = 2.11, and was generated by running the following command on the data in Additional file 3:

~~~
phydms_logoplot results/omegabysite.pdf ‐‐prefs HA_Doud_prefs.csv ‐‐omegabysite
     results/omegabysite.txt ‐‐stringency 2.11 ‐‐minP 0.001
~~~

In this figure, the HA sequence is numbered sequentially beginning with 1 for the first site with deep mutational scanning data, which is the second residue in the protein.

~~~
phydms has vastly superior computational performance to existing alternatives
~~~

Our rationale for developing `phydms` was to enable the analyses described above to be performed faster than with existing software. To validate the improved computational performance, we compared `phydms` (version 2.0.0) to alternative programs that have been used to fit ExpCM. The comparisons used the HA sequences described in Table 1 with ExpCM informed by the deep mutational scanning in [15], and were performed on a single core of a 2.6 GHz Intel Xeon CPU.

Table 5 shows the results. With default settings, `phydms` took 10 minutes to optimize the model parameters and branch lengths. This runtime could be decreased by scaling the branch lengths by a single parameter rather than optimizing them individually (`‐‐brlen scale` option); other work has shown that when the initial tree is reasonably accurate, this approximation can improve runtime while only slightly affecting model fit [47, 48]. Fitting the nucleotide frequency parameters *ϕ*_*w*_(`‐‐fitphi` option) rather than determining them empirically doubled the runtime. The log likelihood and values of the model parameters *β* and *ω* were nearly identical for all three of these settings of `phydms`. Interestingly, the gradient-based optimization appears to be important: using `phydms` without gradients (`‐‐nograd` option) increased the runtime over 5-fold while also yielding a poorer log likelihood.

**Table 5.** Comparison of `phydms` to alternative software for optimizing a tree of 34 HA sequences. HyPhy and `Bio++` use models that fit *ϕ*, whereas by default phydms determines *ϕ*_*w*_ empirically. Log likelihoods are not expected to be identical across software. Full code, data, and results are in Additional file 7.

| software | runtime (min) | log likelihood | $\beta$ | $\omega$ |
| --- | --- | --- | --- | --- |
| phydms, scale branches | 7.8 | -4877.9 | 2.11 | 0.52 |
| phydms, default settings | 10.5 | -4877.7 | 2.11 | 0.52 |
| phydms, fit $\phi$ values | 23.2 | -4876.5 | 2.11 | 0.53 |
| phydms, no gradient | 52.8 | -4894.0 | 2.13 | 0.57 |
| Bio++ via old phydms | 962.6 | -4880.6 | 2.09 | 0.53 |
| HyPhy via phyloExpCM | 2102.0 | -4908.4 | 2.11 | 0.57 |

Two alternative programs have previously been used to fit ExpCM. Early work [22] with these models used a `Python` program `phyloExpCM` (https://github.com/jbloom/phyloExpCM) to run `HyPhy` to optimize ExpCM similar to the ones used here. Subsequent work [23] used an old version of `phydms` to fit ExpCM identical to the ones here using the `Bio++` libraries [25]. We ran both these programs on the HA data set, using `phyloExpCM` version 0.3 with `HyPhy` version 2.22, and `phydms` version 1.3.0 with `Bio++`. Table 5 shows that these programs were ~100-fold and ~200-fold slower than `phydms` version 2.0.0 with default settings. A small portion of the slower runtime is because these earlier implementations cannot calculate empirical nucleotide frequency *ϕ*_*w*_ parameters; however they remain much slower than `phydms` even when these parameters are fit. Divining the reasons for the performance differences was not possible, as the programs differ completely in their implementations. But reassuringly, all programs yielded similar model parameters *β* and *ω* despite independent implementations of the likelihood calculations and the optimization.

## Conclusions

We have described a new software package, `phydms`, that facilitates efficient phylogenetic analysis of gene sequences using substitution models informed by deep mutational scanning experiments. Using these ExpCM, `phydms` can quantitatively compare deep mutational scanning measurements to selection on genes in nature. It can re-scale deep mutational scanning data to account for differences in the stringency of selection between the lab and nature, identify sites evolving differently in nature than expected from the experiments, and compare how well different experiments on the same gene describe natural selection.

The ability to perform these comparisons is useful because the rationale for many deep mutational scanning experiments is to provide information about the effects of mutations on genes in nature. For instance, there are many ways to design an experiment, and it is often not obvious which choices are best if the goal is to make the experiment reflect natural selection – using `phydms`, it is possible to quantitatively compare how well different experiments describe natural selection. Likewise, it is often useful to know which specific sites in a gene are evolving differently in nature than expected from experiments in the lab [23]; `phydms` makes statistically rigorous identification of these sites possible. The speed and ease of use of `phydms` makes these analyses practical for real datasets. As deep mutational scanning data become available for an increasing number of genes, `phydms` will facilitate comparison of the experimental measurements to selection in nature.

## Availability and requirements

- **Project name:** `phydms`
- **Project home page:**

– Documentation: jbloomlab.github.io/phydms/
– Source code: https://github.com/jbloomlab/phydms
- **Operating system(s):** Tested on Linux and Mac OS X
- **Programming language:** `Python`, tested with version 2.7 and version 3.4+
- **Other requirements:** pyvolve [52], `weblogo` [50], and several other widely available `Python` packages.
- **License:** GNU GPLv3
- **Restrictions to use by non-academics:** None

## Abbreviations

ExpCM: experimentally informed codon substitution model
YNGKP: Yang, Nielsen, Goldman, Krabbe-Pedersen substitution model [35]
AIC: Aikake Information Criterion

## Ethics approval and consent to participate

Not applicable.

## Consent for publication

Not applicable.

## Availability of data and material

The latest source code for phydms is available at https://github.com/jbloomlab/phydms. Analyses in this paper used version 2.0.0 of phydms; the code for this version is at https://github.com/jbloomlab/phydms/tree/2.0.0.

## Author’s contributions

SKH and JDB co-wrote the phydms software with contributions from MBD. SKH developed the tutorials. SKH and JDB wrote the initial draft of the manuscript, which all three authors then reviewed and revised.

## Acknowledgements

We thank Erick Matsen, Vladimir Minin, and Joe Felsenstein for helpful comments that aided in the planning and design of the software. We thank Hugh Haddox for assistance in testing the software.

## Additional Files

### Additional file 1

This PDF contains details of the calculations of the likelihood and its gradient as implemented in `phydms`.

### Additional file 2

This ZIP file contains the code, input data, and full results of the `phydms` HA analysis with preferences measured in [15] summarized in Table 2.

### Additional file 3

This ZIP file contains the code, input data, and full results of the stringency parameter comparison with `phydms_logoplot` summarized in Figure 3.

### Additional file 4

This ZIP file contains the code, input data, and full results of the multiple *β*-lactamase deep mutational scan comparison summarized in Table 3.

### Additional file 5

This ZIP file contains the code, input data, and full results of the multiple HA deep mutational scan comparison summarized in Table 4.

### Additional file 6

This ZIP file contains the code, input data, and full results of the phydms ‐‐omegabysite HA analysis with preferences measured in [15] summarized in Figure 4.

### Additional file 7

This ZIP file contains the code, input data, and full results of the program runtime comparison summarized in Table 5.

